# Acute EtOH enhances septohippocampal coordination but disrupts intrinsic hippocampal theta dynamics during foraging

**DOI:** 10.1101/2025.07.22.666144

**Authors:** Caleb B. Darden, Meagan D. Marks, Andrew J. Kesner

## Abstract

Theta oscillations – rhythmic patterns of synchronous activity within discrete brain regions – are known to support memory, navigation, and behavioral coordination, and are sensitive to pharmacological manipulation. Acute ethanol (EtOH) exposure has been shown to alter theta oscillations, but its effects on transient theta bursts and cross-regional coordination during naturalistic behavior remain unclear. We recorded local field potentials (LFPs) from the medial septum (MS), hippocampal *Cornu Ammonis 1* (CA1), and medial prefrontal cortex (mPFC) in freely foraging mice following intraperitoneal injection of EtOH (1.5 g/kg) or saline. We analyzed spectral power, theta burst dynamics, phase, and lag timing. Burst features from CA1 were used to train a machine learning classifier to predict session condition. EtOH impaired locomotion and reduced goal-directed behaviors, particularly early in the session. In CA1, theta power shifted toward lower frequencies and lagged coherence declined. EtOH increased the frequency but reduced the duration of theta bursts in CA1, and in MS, only burst count increased. EtOH enhanced the temporal alignment of MS–CA1 burst pairs. Phase-locking between CA1 and MS during coupled bursts remained present but showed altered structure. Our classifier achieved robust performance using burst features such as skew and entropy, and reliably distinguished treatment conditions. EtOH modulates septohippocampal dynamics by altering the timing and structure of theta bursts. These results suggest that burst-level features are sensitive markers of EtOH’s circuit-level effects during naturalistic behavior.

## Introduction

To survive animals must appropriately sense and respond to environmental stimuli. The brain is organized in a manner to coordinate these processes at many levels, from the activity of single neurons to macroscale network activity and communication between various brain regions. Neuronal function within the brain is predominantly studied through the lens of single unit activity, i.e. neuronal action potentials, or “spikes”.

However, this spike-centric approach does not typically address electric fields that arise from the coordinated transmembrane currents of neuronal ensembles and can modulate excitability and timing on millisecond timescales [1]. Experimental and computational studies show that weak electric fields, as low as 0.2–0.5 V/m, can reliably shift spike timing, synchronize neural populations, and entrain network dynamics at physiologically relevant field strengths [2–4]. These effects are particularly relevant in spatially organized structures like the hippocampus, where aligned pyramidal cell dendrites enhance sensitivity to field effects [2, 5].

Furthermore, oscillating electric fields may amplify these effects through resonance with intrinsic membrane and network properties, creating temporally structured ‘windows of excitability’ that shape how ensembles encode, transmit, and receive information [6–8]. Within this framework, LFPs and their underlying electric fields are functionally active components of neural computation, coordinating large-scale interactions via rhythmic, sub-threshold mechanisms.

Among these rhythmic patterns, theta oscillations (4–12 Hz) are prominent in the mammalian brain and fundamentally coordinate neural activity across distributed circuits [9]. Crucially involved in memory formation, spatial navigation, and behavioral coordination, hippocampal theta oscillations can be generated intrinsically or driven by medial septum (MS) GABAergic input [10–12]. MS GABAergic neurons exhibit phasic firing aligning to CA1 theta, primarily via projections to hippocampal interneurons that support theta generation [13, 14].

Theta oscillations’ influence extends beyond the hippocampus or septohippocampal circuit and establishes coherent networks with the mPFC and other regions. Simultaneous recordings from the hippocampus and mPFC reveal enhanced theta-frequency synchrony that is specifically recruited during spatial working memory tasks [15]. This hippocampal-prefrontal theta coherence mediates working memory performance, with synchrony strength correlating with behavioral accuracy [16, 17]. Theta oscillations coordinate interactions between these regions during complex cognitive operations, and hippocampal theta has been shown to modulate mPFC gamma oscillations to facilitate interregional communication [18]. Hippocampal theta’s functional role extends beyond network synchrony to powerful modulation of learning. For example, in an eyeblink classical conditioning study, training during hippocampal theta periods dramatically accelerated associative learning, with animals acquiring the conditioned response in half the trials of non-theta periods [19].

Thus, the septohippocampal pathway presents an ideal system to investigate how field-based communication structures brain dynamics. Recent studies reveal that cortical gamma and beta oscillations appear as discrete bursts in LFP recordings and that these high-amplitude events support working memory processes [20]. However, it is unknown whether theta-band activity in the septohippocampal circuit is similarly organized.

Despite exploration of LFP-LFP dynamics in some corticohippocampal circuits, the existence and role of burst-like LFP events in rodent theta have received limited attention. Human studies, however, report transient theta bursts under specific conditions. For example, during transcendental meditation, theta bursts of approximately 1.8 seconds recur every two minutes [21], and in the human hippocampus during virtual navigation, theta appears as discrete bursts throughout movement [22]. These findings suggest an event-based, rather than continuous, mode of theta operation and raises the possibility that theta bursts in rodents serve a similar functional role.

EtOH profoundly modulates multiple neurotransmitter systems that regulate theta oscillations, including GABAergic, glutamatergic, and cholinergic networks [23]. Acute EtOH exposure produces rapid changes in GABAA receptor function, subunit composition, thereby affecting inhibitory signaling [24]. Prior studies demonstrate that acute EtOH disrupts theta oscillations, leading to attenuated hippocampal theta rhythm, dampened spontaneous firing, and strengthened GABA-mediated inhibition in MS neurons [25–27]. However, no research has explored the effects of EtOH or other theta- altering compounds on LFP-LFP interregional communication, let alone burst dynamics, in rodents.

The present study addresses these knowledge gaps, investigating whether theta-band LFP activity exhibits functionally meaningful burst dynamics in the mouse brain and how these features are disrupted by acute EtOH exposure. We employed simultaneous local field potential recordings from MS, CA1, and mPFC in freely behaving mice during a foraging task (Fig. 1A-C). We hypothesized that theta bursts would occur in the MS and CA1, display some form of temporal entrainment, and that EtOH would alter this coordination, manifesting as altered burst characteristics, disrupted temporal relationships, and reduced directional coupling between these regions. Understanding these mechanisms at the level of individual oscillatory events provides new insights into how the mammalian brain communicates, and how acute EtOH intoxication compromises its function.

**Figure 1.**
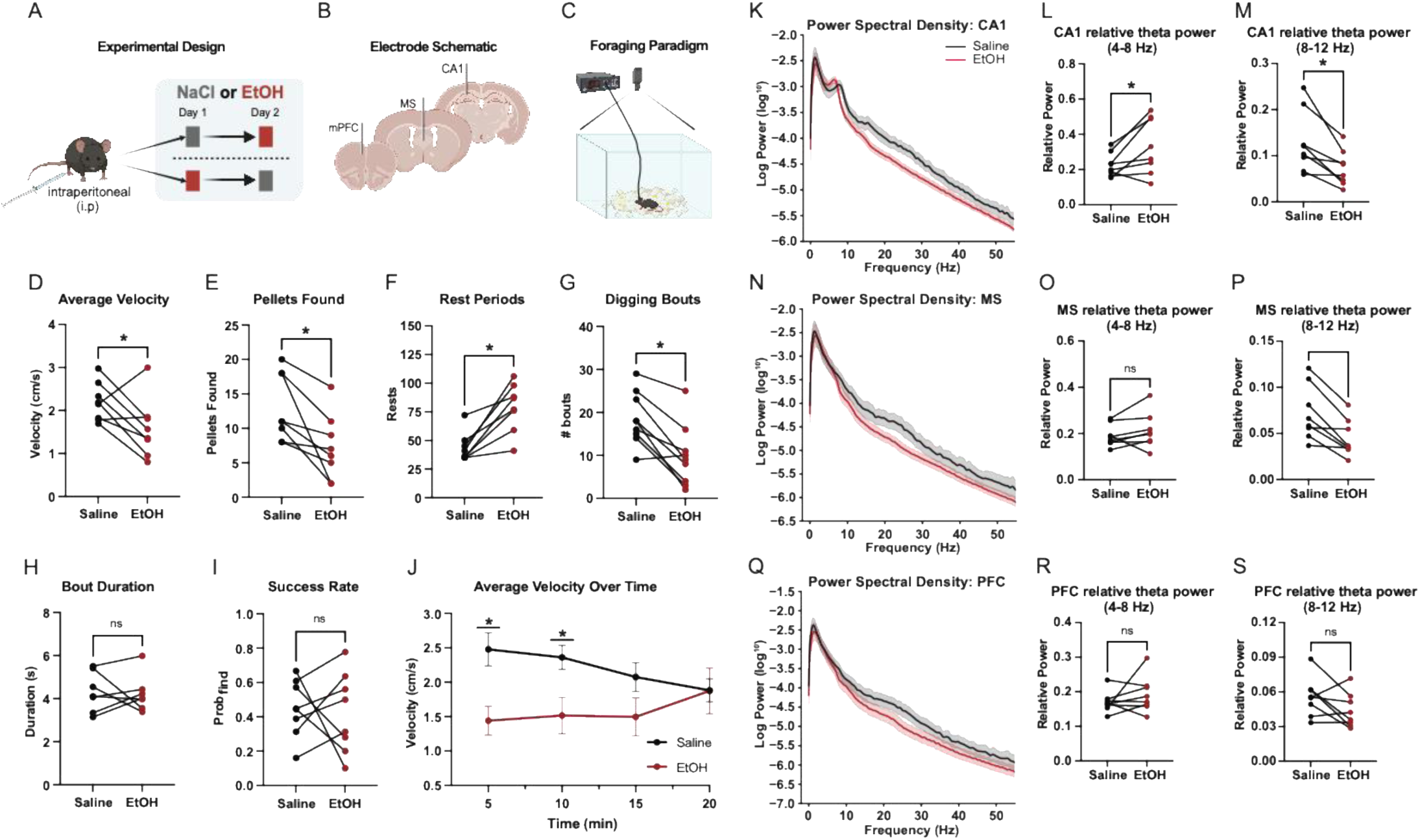
Acute EtOH exposure disrupts attenuates locomotion and Theta power. (A) Schematic of our within-subject, counterbalanced administration of 1.5 g/kg EtOH (red) and saline (gray). (B) Unilateral microwire implant sites in mPFC, MS, and dorsal CA1. (C) Overview of foraging paradigm with top-down video recording. (D) Average velocity during free foraging following EtOH administration compared to saline (paired t-test, *p* = 0.0487). (E) Food pellets recovered during foraging task (paired t-test, *p* = 0.0113). (F) Rest period count (defined as velocity < 0.5 cm/s for ≥2 s) during foraging task (paired t-test, *p* = 0.0018). (G) Digging bout counts by treatment type (paired t-test, *p* = 0.0011) (H) Digging bout count duration by treatment type (paired t-test, *p* = 0.8528). (I) Proportion of pellet yielding digging bouts by treatment type (*p* = 0.7921). (J) Group average velocity across foraging session (two-way ANOVA | Post hoc Holm– Sidak tests | 5 min (*p* = 0.0142, Δ = 1.036 cm/s); 10 min (*p* = 0.0470, Δ = 0.8463 cm/s); 15 min: *p* = 0.1802; 20 min: *p* = 0.9812). Data are shown as mean ± SEM. (K) Power spectral density curves in CA1 by treatment type (grouped) across trimmed frequency spectrum (0 – 55Hz). (L) Relative low-theta power in CA1 by treatment type (paired t-test, *p* = 0.0396). (M) Relative high-theta power in CA1 by treatment type (paired t-test, *p* = 0.0084). (N) Power spectral density curves in MS by treatment type (grouped) across trimmed frequency spectrum (0 – 55Hz). (O) Relative low-theta power in MS by treatment type (paired t-test, *p* = 0.2545). (P) Relative high-theta power in MS by treatment type (paired t-test, *p* = 0.0029). (Q) Power spectral density curves in mPFC by treatment type (grouped) across trimmed frequency spectrum (0 – 55Hz). (R) Relative low-theta power in mPFC by treatment type (paired t-test, *p* = 0.3179). (S) Relative high-theta power in mPFC by treatment type (paired t-test, *p* = 0.1120).

## Methods and Materials

### Subjects

Eight adult, 4mo, vGlut2-Cre mice (4 male, 4 female) were used. All animals were maintained on a 12:12 h light/dark cycle and food-restricted to 90% of baseline body weight during behavioral testing. All methods used in this work were approved by the Animal Care and Use Committee of the National Institute on Alcohol Abuse and Alcoholism (protocol #: LIN-DL-1) and were within the guidelines described in the NIH Guide to the Care and Use of Laboratory Animals. Of note, mice in this experiment were prepared for another set of experiments involving chemogenetic modulation of MS glutamate neuron during goal-directed tasks, but these studies are not presented in this manuscript, nor were any chemogenetic perturbations performed. See supplementary information for a full description of subjects.

### Surgical Procedures

Under isoflurane anesthesia, mice were implanted with custom-built microwire electrodes for chronic in vivo LFP recording. Each electrode consisted of a single PFA-coated tungsten wire (0.005 “bare, 0.007” coated; AM Systems), stripped at one end, inserted into a gold pin, and implanted manually. One electrode was targeted to each of the following regions: MS, CA1, and mPFC. A reference electrode (0.008“ bare, 0.010” coated) was implanted over the occipital plate. Dura was removed using a fine needle to reduce insertion resistance. Implants were secured with dental cement into a 3D-printed head cap. Mice recovered for one week with ad libitum access to food and water.

Stereotaxic coordinates were as follows (relative to bregma; AP, ML, DV): MS: +0.8, 0.0, -3.3; CA1: -2.3, +2.0, -1.4; mPFC: +1.7, +0.4, -2.0. Refer to supplementary methods (Supp Fig. 2) for electrode placements. In the same surgical procedure, animals also received microinjections of AAV2-hSyn-DIO-mCherry, -Gq, or -Gi vectors targeting glutamatergic neurons in the medial septum, however, again, no chemogenetic manipulations were performed in the present study.

**Figure 2.**
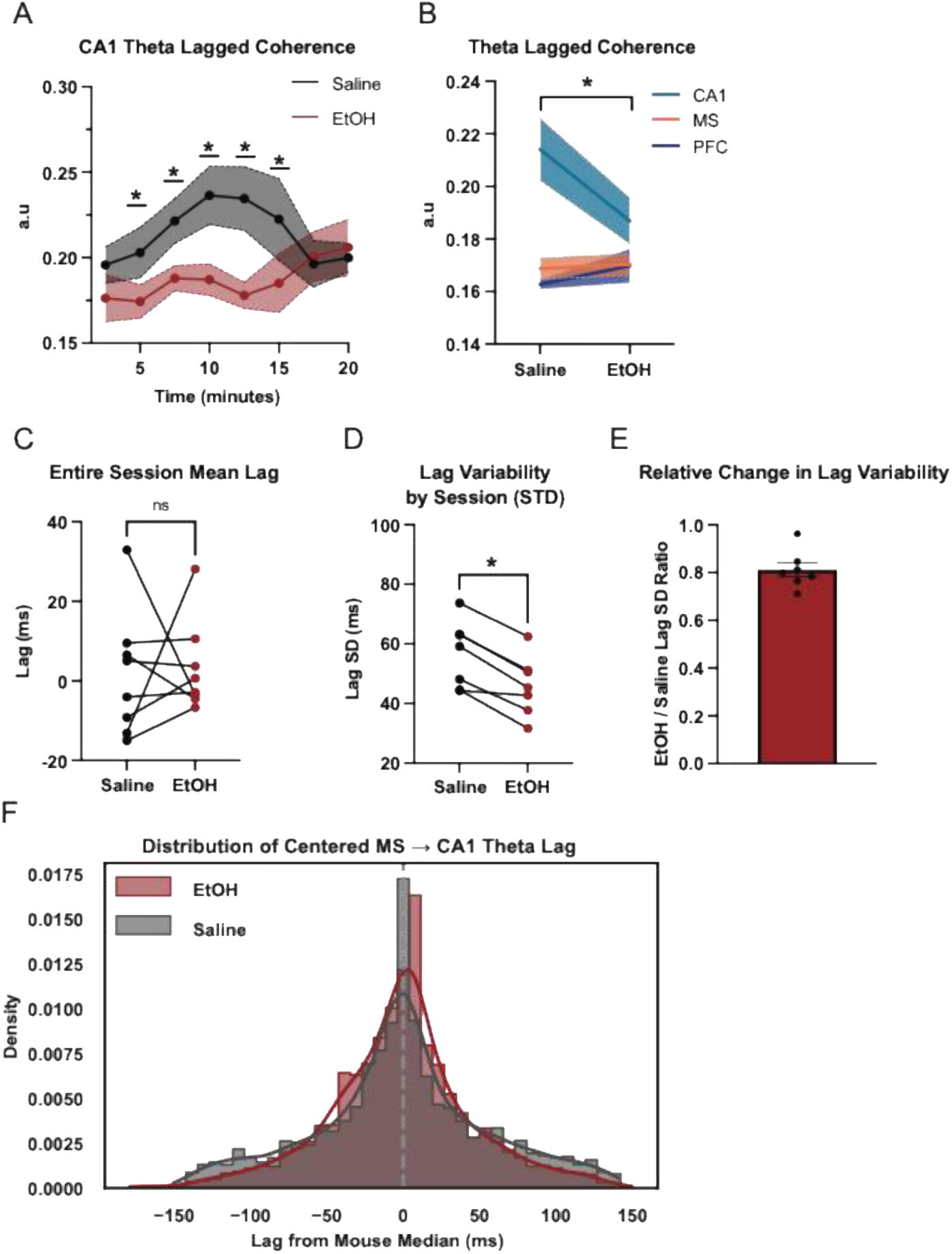
EtOH reduces CA1 theta rhythmicity as septohippocampal theta stability. (A) Time-resolved analysis of CA1 theta lagged coherence (two-way ANOVA, Time: F = 1.204, p = 0.3183; Treatment: F = 5.323, p = 0.0544; Interaction: F = 2.834, p = 0.0146 | Post hoc Holm–Šídák tests: 5 min (p = 0.0401), 7.5 min (p = 0.0173), 10 min (p = 0.0007), 12.5 min (p = 0.0001), and 15 min (p = 0.0081). (B) Session-averaged theta lagged coherence for CA1, MS, and mPFC (paired t-test: CA1, p = 0.0435; MS, p = 0.7733; mPFC, 0.3663]. Shaded areas indicate SEM. (C) Representative MS→CA1 theta lag values by treatment type (paired t-test, p > 0.05). (D) Within-mouse standard deviation of MS→CA1 theta lag values (paired t-test, p < 0.001). (E) Bar plot showing EtOH/saline ratio for MS→CA1 theta lag SD values. (F) Kernel density estimates of MS→CA1 theta-lag distributions, with each mouse’s session lags centered by subtracting its median.

### Behavioral Paradigm

Mice were habituated to the foraging arena over five days and to food pellets in the home cage for three days. During test sessions, animals foraged freely in an 8×13 inch arena within a sound-attenuating chamber for 20 minutes. Each session contained 20 food pellets, with 12 partially buried to encourage exploratory digging, i.e. foraging for food. Mice received either an intraperitoneal (i.p.) saline injection or 1.5 g/kg EtOH (20% v/v in saline) 3 minutes before the session. Sessions were counterbalanced across days within subjects. Prior to the foraging paradigm, all animals had previously undergone behavioral testing with chemogenetic approaches. However, roughly 3 weeks had passed with no chemogenetic or behavioral assays performed prior to the currently reported experiment.

### Video Tracking

Behavior was recorded using an overhead USB camera at 20 FPS. EZTrack (Pennington, Z. T et al., 2019) was used to extract 2D position, velocity (cm/s), and acceleration (cm/s²). Frames were synchronized with LFP recordings. Event-related behavioral metrics (digging bouts, durations of bouts, and whether a reward pellet was found for a respective bout) were manually scored from video by a researcher blinded to experimental conditions.

### LFP Acquisition and Filtering

LFPs were recorded using a Tucker-Davis Technologies PZ5 amplifier at a 1017 Hz sampling rate using Intan head stages. Signals were notch filtered at 60 Hz to remove line noise and then bandpass filtered into standard frequency bands using zero-phase finite impulse response (FIR) filters: Delta (0.5–4 Hz), Theta (5–12 Hz), Beta (13–20 Hz), Low Gamma (30–55 Hz), and High Gamma (65–100 Hz). All filters were implemented using custom scripts designed to ensure narrow transition bands, minimal phase distortion, and high stopband attenuation. Power spectral density (PSD) was estimated using Welch’s method with 6-second non-overlapping windows (nfft = 8096). Relative band power was calculated by dividing the power in each band by the total power between 0–55 Hz. This allowed normalization across animals and conditions for comparison of oscillatory activity.

### LFP postprocessing and Analysis

We performed phase and lag analysis using established methods. See supplemental methods for full description of these analysis.

#### Phase Analysis

To examine EtOH’s impact on septohippocampal LFP-LFP timing, we computed the instantaneous phase offset between MS and CA1 within the theta band. For analyses in which preferred phase angle offset was used, phase differences were circularly averaged across time and summarized using kernel density estimates (KDEs) per mouse. Group comparisons (saline vs. EtOH, coupled vs. uncoupled bursts) were made using permutation tests and Watson-Williams circular ANOVA. For cross-region coordination, we calculated peri-burst PLVs between CA1 and MS. Theta bursts in CA1 were classified as “coupled” if preceded by an MS theta burst within 400 ms. For each coupled event, PLV was calculated from 2-second pre- and post-burst windows. PLV was defined as the magnitude of the mean vector of phase differences between Hilbert-transformed CA1 and mPFC signals. Post/pre PLV ratios were computed for each burst and averaged per mouse for statistical comparison. See supplementary methods for a full description of phase analysis.

#### Lag Analysis

To evaluate the directionality and temporal consistency of MS–CA1 coordination, we estimated lags between theta amplitude envelopes using cross-correlation [28].

MS→CA1 delays were computed in overlapping windows, and lag variability was compared across saline and EtOH sessions. See supplementary methods for a full description of lag analysis.

#### Burst Detection

CA1 LFPs were bandpass filtered (5–12 Hz) using zero-phase Butterworth filters and segmented into non-overlapping 5-second windows. A dual-threshold algorithm applied to these bandpass-filtered signals [29]. For downstream analyses, bursts were further filtered to exceed a minimum duration of 1.5 cycles of the band’s low cutoff frequency (e.g., ≥200 ms for theta). See supplementary methods (Sup Fig. 1) for full description of burst detection algorithm.

#### Machine Learning Classification

To determine whether CA1 theta bursts encoded treatment condition, we trained an XGBoost classifier on burst features. Bursts were labeled by condition and characterized using waveform, spectral, and contextual metrics. Classification accuracy was evaluated using ROC and precision–recall curves. See supplementary methods for a full description of the classification pipeline.

#### Statistics and Visualization

All statistical analyses were performed in Python using custom scripts built with SciPy, NumPy, NeuroDSP, and statsmodels, or in GraphPad Prism (version 11; GraphPad Software, La Jolla, CA, USA). Figures were generated using Seaborn, Matplotlib, and GraphPad Prism. All data and analysis scripts are available on GitHub (to be linked upon peer review publication). All statistical tests and results are reported in figure legends.

## Results

### EtOH diminishes locomotion and foraging task performance

Acute EtOH administration at a dose known to reduce locomotor output without full sedation [30], indeed impaired both locomotion and foraging behavior in mice during free exploration. Relative to saline sessions, EtOH reduced average velocity (Fig. 1D) and decreased the number of food pellets discovered (Fig. 1E). EtOH also increased the number of rest periods (Fig. 1F) and reduced the total number of digging bouts (Fig. 1G), though the average duration of each bout (Fig. 1H) and the proportion of successful digs (Fig. 1F) remained unchanged. EtOH-induced reduction in velocity was most evident early in the session, with differences at 5 and 10 minutes but not at later time points (Fig. 1J). These results indicate that EtOH acutely suppresses task engagement, particularly in the initial phase of behavior.

### EtOH alters theta power in MS and CA1

EtOH administration led to region-specific changes in theta-band activity across the hippocampal–septal–prefrontal circuit. In CA1, power spectral density (PSD) curves revealed a shift in theta dynamics, with increased power in the low-theta range (4–8 Hz) and decreased power in the high-theta range (8–12 Hz) during EtOH sessions (Fig. 1K). This shift was reflected in a significant increase in relative low-theta power and a corresponding decrease in high-theta power (Fig. 1L–M). In the MS, EtOH broadly suppressed theta activity, with PSD curves showing a general reduction across the theta band (Fig. 1N). While low-theta power in MS remained unchanged (Fig. 1O), high-theta power was significantly reduced (Fig. 1P). In contrast, mPFC theta activity appeared relatively unaffected, with minimal changes in PSD profiles (Fig. 1Q) and no significant differences in either low- or high-theta power (Fig. 1R–S). These findings suggest that EtOH selectively alters theta-band dynamics in CA1 and MS, with a pronounced reduction in high-theta oscillations.

### EtOH Disrupts Hippocampal Theta Rhythmicity

To assess the temporal stability of theta oscillations, we measured lagged coherence over the course of the foraging session. Lagged coherence is a measure of how consistent the phase is of an oscillation over time (Fransen et al., 2015). In CA1, EtOH administration led to a progressive disruption of the stability of theta rhythmicity, with significantly reduced lagged coherence between 5- and 15-minutes post-injection relative to saline (Fig. 2A). This effect appeared specific to CA1, as session-averaged lagged coherence across brain regions revealed no significant change in MS or mPFC (Fig. 2B). These findings suggest that EtOH selectively impairs sustained theta rhythmicity in the hippocampus, particularly during the early to mid-portions of this foraging task.

### EtOH reduced MS-CA1 theta lag variability but not overall timing

To evaluate how EtOH affects the temporal consistency of directional septohippocampal coordination, we analyzed MS→CA1 theta lags using amplitude envelope cross-correlation. Absolute lag values varied by mouse, lacked consistent directionality, and these metrics were not meaningfully altered by EtOH (Fig. 2C). To account for inter-animal differences in absolute lag values, distributions were centered by subtracting each mouse’s session-specific median lag. While the shape of centered lag distributions remained largely similar between conditions, EtOH significantly decreased within-mouse lag variability as shown by the standard deviation of a mouses absolute lag value distribution across the session (Fig. 2, D-F), EtOH alters the consistency of septohippocampal communication by constraining the temporal variability of MS→CA1 theta transmission. This suggests that while the directionality of coordination remains unchanged, EtOH may restrict the dynamic flexibility of this pathway, potentially reducing the system’s capacity to adaptively shift timing relationships during behavior.

### EtOH Alters Theta Burst Dynamics in CA1 and MS

Given the results from our power analyses, we expected that, if we did observe transient oscillatory events in a frequency band, dynamics in CA1 theta would be most responsive to EtOH administration. Nonetheless, we deployed the algorithm to detect bursts in all regions and across frequency bands. Indeed, theta burst dynamics were most strongly altered by EtOH. In CA1, EtOH increased the total number of theta and beta bursts (Fig. 3A) and significantly shortened average theta burst duration (Fig. 3B). Time-resolved analyses revealed elevated theta burst counts during the latter half of the session (10–20 min; Fig. 3C) and reduced burst duration in the middle portion of the session (5–15 min; Fig. 3D). EtOH also shifted the distribution of CA1 burst durations toward shorter events (Fig. 3E). In the MS, EtOH similarly increased theta and beta burst counts (Fig. 3F) but did not affect overall burst duration (Fig. 3G). A time-by-treatment interaction showed that burst counts were elevated in the final time bin of the session (Fig. 3H), while duration remained stable across time (Fig. 3I), with no shift in duration distribution (Fig. 3J). In contrast, mPFC showed no overall effect of EtOH on burst count or duration (Fig. 3K–L). However, time-resolved analysis revealed transient increases in burst count at the start and end of the session (Fig. 3M), with no accompanying change in duration (Fig. 3N) or distribution (Fig. 3O). These results suggest that EtOH enhances the occurrence of theta bursts in CA1 and MS, with a selective reduction in burst duration confined to CA1.

**Figure 3.**
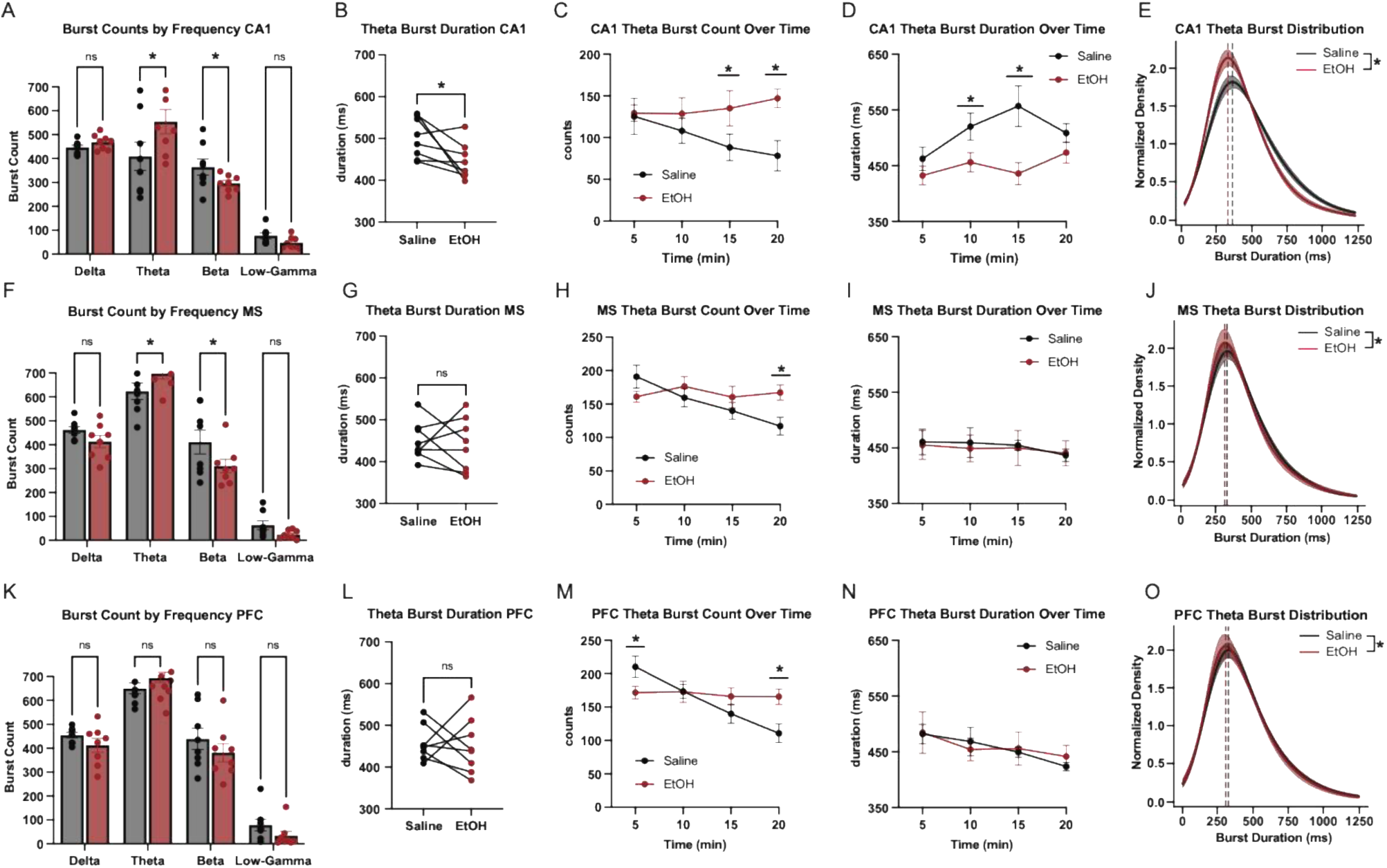
Both CA1 and MS show theta burst activity, and their architecture is modulated by EtOH. (A) Burst count in CA1 across frequency bands [paired t-test, theta, p < 0.0001; beta, p = 0.0168]. (B) Average duration of CA1 theta bursts (*paired t-test, p = 0.0370)*. (C) Time-resolved analysis of CA1 theta burst count (two-way ANOVA, Time: F = 1.534, p = 0.5762; Treatment: F = 12.06, p = 0.0025; Interaction: F = 6.078, p = 0.040 | Post hoc Tukey tests: 10–15 min (p = 0.0073); 15–20 min (p = 0.0003). (D) Time-resolved analysis of CA1 theta burst duration (two-way ANOVA, Time: F = 18.65, p = 0.0002; Treatment: F = 4.39, p = 0.0373; Interaction: F = 2.77, p = 0.0438 | Post hoc Tukey tests: 5–10 min p = 0.0357; 10–15 min p = 0.0003). (E) Normalized KDE showing CA1 theta burst duration distributions by treatment type (*Kolmogorov–Smirnov test*, D = 0.077, p *< 0.0001*). (F) Burst count in MS across frequency bands [paired t-test, theta, p = 0.0306; beta, p = 0.0033]. (G) Average duration of MS theta bursts (paired t-test, p = 0.6823). (H) Time-resolved analysis of MS theta burst count (two-way ANOVA, Time: F = 2.771, p = 0.0669; Treatment: F = 9.892, p = 0.0163; Interaction: F = 3.493, p = 0.0337 | Post hoc Tukey tests: 15–20 min (p = 0.0100). (I) Time-resolved analysis of MS theta burst duration (two-way ANOVA, Time: F = 0.6843, p = 0.5716; Treatment: F = 0.0303, p = 0.86667; Interaction: F =0.1196, p = 0.9475 | Post hoc Tukey tests: not significant at any timepoints. (J) Normalized KDE showing MS theta burst duration distributions by treatment type (*Kolmogorov–Smirnov test*, D = 0.043, p = 0.0001) (K) Burst count in mPFC across frequency bands [not significant]. (L) Average duration of mPFC theta bursts (paired t-test, p = 0.8124). (M) Time-resolved analysis of mPFC theta burst count (two-way ANOVA, Time: F = 4.473, p = 0.0140; Treatment: F = 4.597, p = 0.0692; Interaction: F = 5.393, p = 0.0065 | Post hoc Tukey tests: 0-5 min (p = 0.0346), 15–20 min (p = 0.0043). (N) Time-resolved analysis of mPFC theta burst duration (two-way ANOVA, Time: F = 3.341, p = 0.0388; Treatment: F = 0.01217, p = 0.9153; Interaction: F = 0.6617, p = 0.5848 | Post hoc Tukey tests: not significant at any timepoints. (O) Normalized KDE showing MS theta burst duration distributions by treatment type (*Kolmogorov–Smirnov test*, D = 0.034, p = 0.0045).

### EtOH enhances temporal alignment of MS-CA1 Bursts

To examine how EtOH influences the timing of septohippocampal coordination, we computed the delay between theta bursts in MS and the next theta burst in CA1. Subsequent CA1 bursts that did not occur within 1 second of MS theta burst completion were excluded. By generating kernel density estimates (KDE) of these MS→CA1 burst latencies, we assessed whether EtOH alters the distribution of interregional burst propagation delays (Fig. 4A). To examine how the temporal structure of MS→CA1 interactions relates to local burst dynamics, we analyzed the relationship between MS and CA1 theta burst durations as a function of their temporal separation. Using the 25th and 75th percentiles of the empirical delay distribution, burst pairs were categorized into short- and long-latency groups. We then compared burst duration correlations across saline and EtOH conditions to assess whether EtOH alters inter-regional coordination as a function of delay. The presence of a stronger correlation between MS and CA1 burst duration under EtOH, particularly in short-latency pairs (Fig. 4B), suggests that brief MS→CA1 delays are associated with tighter temporal coupling and more similar burst structure. In contrast, long-delay pairs showed weaker and less consistent duration coupling (Fig. 4C), regardless of condition, suggesting that extended propagation times may decouple local burst dynamics from upstream temporal structure.

**Figure 4.**
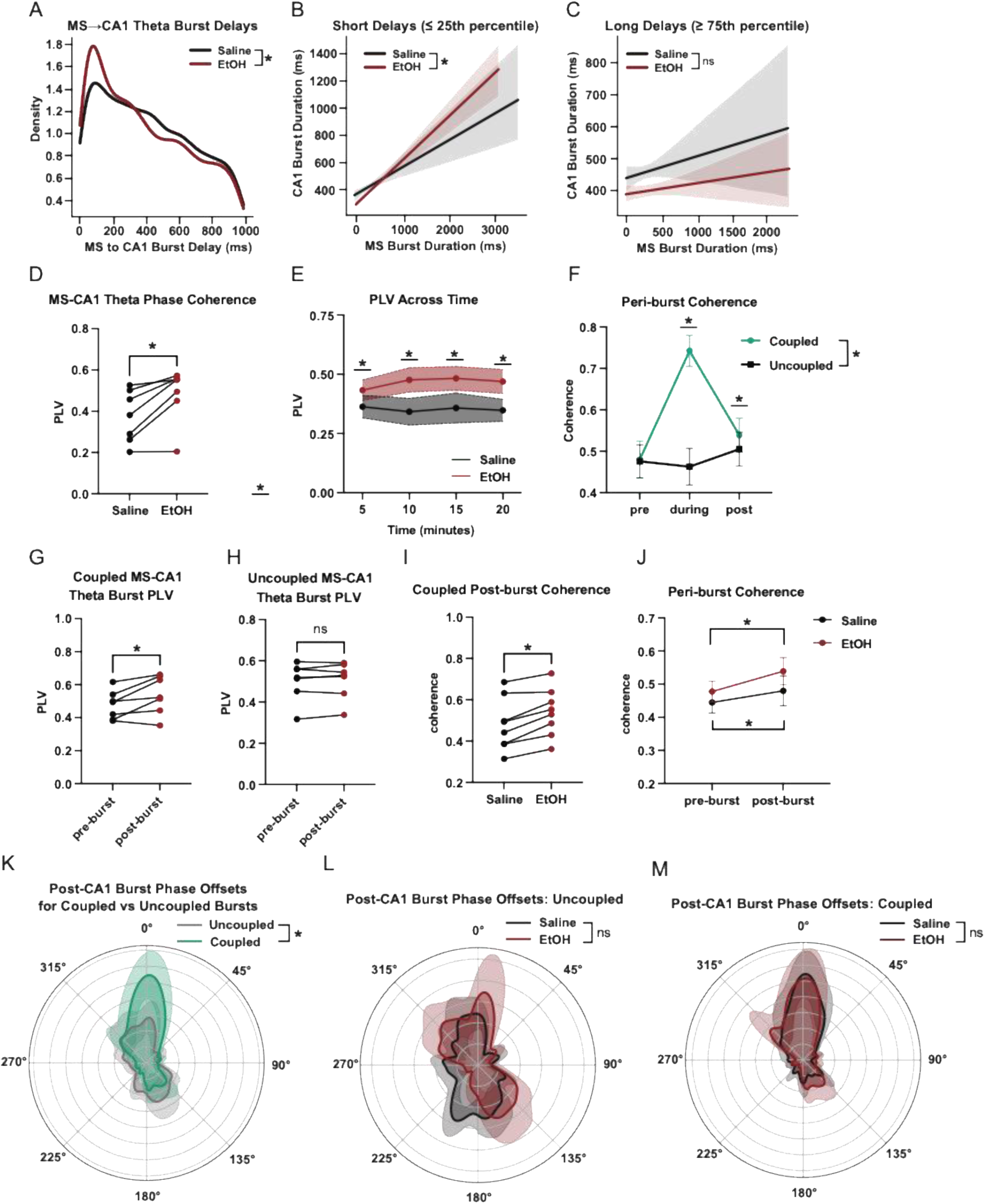
EtOH alters the temporal alignment and interregional coupling of MS→CA1 theta bursts. (A) Kernel density estimates of MS→CA1 theta burst delays (Mann–Whitney U test: U = 2663901.5, p = 1.0484e-03). (B) Short-delay MS→CA1 theta burst pairs for Saline (grey) and EtOH (Red) (EtOH: r = 0.443, n = 1022; saline: r = 0.257, n = 551; Fisher z = –4.01, p = 0.0001). Shaded regions indicate 95% confidence intervals on the regression fit. (C) Long-delay MS→CA1 theta burst pairs for Saline (grey) and EtOH (Red) (EtOH: r = 0.037, n = 879; Saline: r = 0.069, n = 686; Fisher z = 0.62, p = 0.5353). Shaded regions indicate 95% confidence intervals on the regression fit. (D) MS-CA1 session-wide theta PLV (paired t-test, p = 0.0116). (E) Time-resolved MS-CA1 theta PLV across the foraging session (two-way ANOVA, Time: F = 0.3250, p = 0.8073; Treatment: F = 17.76, p = 0.0040; Interaction: F = 0.9998, p = 4124 | Post hoc Holm–Šídák tests: 0-5 min (p = 0.0270), 5-10 min (p = 0.0.0002), 10-15min (p = 0.00031), 15–20 min (p = 0.0043). (F) Coupled (green) versus uncoupled (black) peri-burst MS-CA1 theta coherence showing pre-burst, during burst, and post-burst values (two-way ANOVA, Time: F = 146.1, p < 0.0001; Burst-type: F = 144.0, p = < 0.0001, Interaction: F = 145.7, p < 0.0001 | uncorrected Fisher’s LSD: during-burst (p < 0.0001), post-burst (p = 0.0175). (G) Coupled MS→CA1 pre/post PLV (post-burst: red, pre-burst: black) (paired t-test, p = 0.0012). Pre-burst represents 2 seconds before MS burst onset and post-burst represents 2 seconds after CA1 burst offset. (H) Uncoupled MS→CA1 peri-burst PLV (2 seconds before MS burst onset to 2 seconds after CA1 burst offset) by treatment type (post-burst: red, pre-burst: black) (paired t-test, p = 0.4671). (I) Coupled MS→CA1 post-burst coherence by treatment type (EtOH: red, Saline: black) (paired t-test, p = 0.0032). (J) MS→CA1 peri-burst theta coherence by treatment type (EtOH: red, Saline: black) (paired t-test: Saline pre/post, p = 0.0019; EtOH pre/post, p = < 0.0001). (K) Polar phase distributions of MS–CA1 theta phase offsets after coupled (green) and uncoupled (grey) bursts (Rayleigh test: saline, p < 0.0001, Z = 15109.34; EtOH, p < 0.0001, Z = 31466.76 | Mardia–Watson–Wheeler (MWW), W = 144775.031, p < 0.0001). (L) Polar phase distributions of MS–CA1 theta phase offsets after uncoupled bursts by treatment type (EtOH: red, Saline: black) (Rayleigh test: Saline, p = 0.00015, Z = 8.75; EtOH, p = 0.0085, Z = 6.21 | MWW, W = 14.759, p = 0.3341). (M) Polar phase distributions of MS–CA1 theta phase offsets after coupled bursts by treatment type (EtOH: red, Saline: black) (Rayleigh test: Saline, p < 0.0001, Z = 140.61; EtOH, p < 0.0001, Z = 201.66 | MWW, W = 11.386, p = 0.4115).

### Session-wide and peri-burst phase metrics are altered by EtOH

To assess the degree of oscillatory coherence between the MS and CA1, we computed per mouse/session PLVs as described in the methods. Values from each 8-second window were averaged across time to yield a PLV representative of the session. EtOH increased these MS-CA1 values (Fig. 4D). To assess PLV metrics across time, we used averages from the previously calculated 8 second windows, averaged each 5 minutes, and found that EtOH elevated PLVs throughout the recording session (Fig. 4E).

To learn how EtOH affects peri-burst phase alignment between the MS and CA1, we analyzed theta-band phase offsets in the 1.25-second window following CA1 theta bursts that were preceded by an MS theta burst within 200 ms. For each such MS→CA1 burst pair, we computed the instantaneous phase of both signals in the 5–12 Hz band using the Hilbert transform and calculated the circular difference between them. Each instance yielded a single mean phase offset, representing the between MS and CA1 during the post-burst interval. To determine how theta bursts contribute to cross-regional synchrony, we examined peri-burst PLVs between MS and CA1.

Coupled burst pairs—defined as MS–CA1 theta bursts occurring within 1 second of each other—were associated with markedly higher PLV values than uncoupled events during the post-burst period (Fig. 4G–H), indicating stronger transient synchrony during tightly timed burst interactions. For each event, a single mean phase offset was computed from the Hilbert-transformed theta signal, capturing the directional phase relationship between MS and CA1 throughout the post-burst interval. This phase-based coordination was notably stronger when bursts were temporally aligned, suggesting that close temporal proximity promotes interregional phase coherence. EtOH further enhanced this effect, selectively increasing PLV following coupled bursts (Fig. 8I). This pattern indicates that EtOH may amplify phase-locking during specific epochs of tightly coupled interregional activity. Time-resolved analyses confirmed that this increase in coherence was restricted to the post-burst window, with no comparable elevation in the pre-burst period. A two-way ANOVA revealed that both time and condition contributed to post-burst PLV differences, with EtOH specifically elevating synchrony after closely timed bursts (Fig. 4I).

To visualize the structure of these offsets, we estimated circular phase distributions across mice and conditions. Individual histograms of phase offsets were computed for each mouse within each condition, binned into 360 angular segments from –π to π. To determine whether the observed phase offsets were non-uniform (i.e., whether they exhibited significant directional concentration), we applied the Rayleigh test for circular uniformity. This test quantifies whether phase values are randomly distributed or cluster around a preferred direction; significant results indicate meaningful phase locking.

Rayleigh tests revealed robust non-uniformity in the coupled group under both saline and EtOH conditions (Fig. 4M), and weaker or absent locking in uncoupled events (Fig. 4L).

Finally, to test whether temporal coupling between bursts influenced phase relationships, we compared phase offset distributions between “coupled” (≤1 s) and “uncoupled” MS–CA1 burst pairs (Fig. 4K). This analysis revealed a significant difference between the two groups, indicating that tightly coordinated MS→CA1 theta bursts are associated with stronger and more consistent theta phase alignment during the post-burst period.

### CA1 Theta Bursts relate to locomotor dynamics

To evaluate how hippocampal theta bursts relate to spontaneous motor initiation and movement vigor, we analyzed LFP and locomotor behavioral data under saline and EtOH conditions. Theta power positively correlated with velocity with a significantly stronger correlation and relationship in EtOH compared to saline sessions (Fig. 5A). To determine whether theta bursts are temporally aligned to movement initiation, we first identified bouts of spontaneous motor onset—defined as a transition from at least 2 seconds of rest (velocity < 0.5 cm/s) to sustained movement (velocity > 1.5 cm/s). EtOH treated animals showed an increase in the number of these initiations (Fig. 5B). To determine if these movement bouts were related to theta bursts, quantified the presence and type of CA1 theta bursts surrounding movement initiations (+/- 2s). CA1 theta bursts were observed in this time window for both saline and EtOH sessions, with a greater percentage observed during EtOH (Fig. 5C). To determine when in this window bursts were occurring, we constructed histograms of burst times within the window around movement onset, smoothed them using a Gaussian kernel (σ = 1 bin), normalized to unit area, and averaged across mice. Burst density peaked immediately before movement onset in both conditions with no difference in the temporal distribution observed between groups (Fig. 5D).

**Figure 5.**
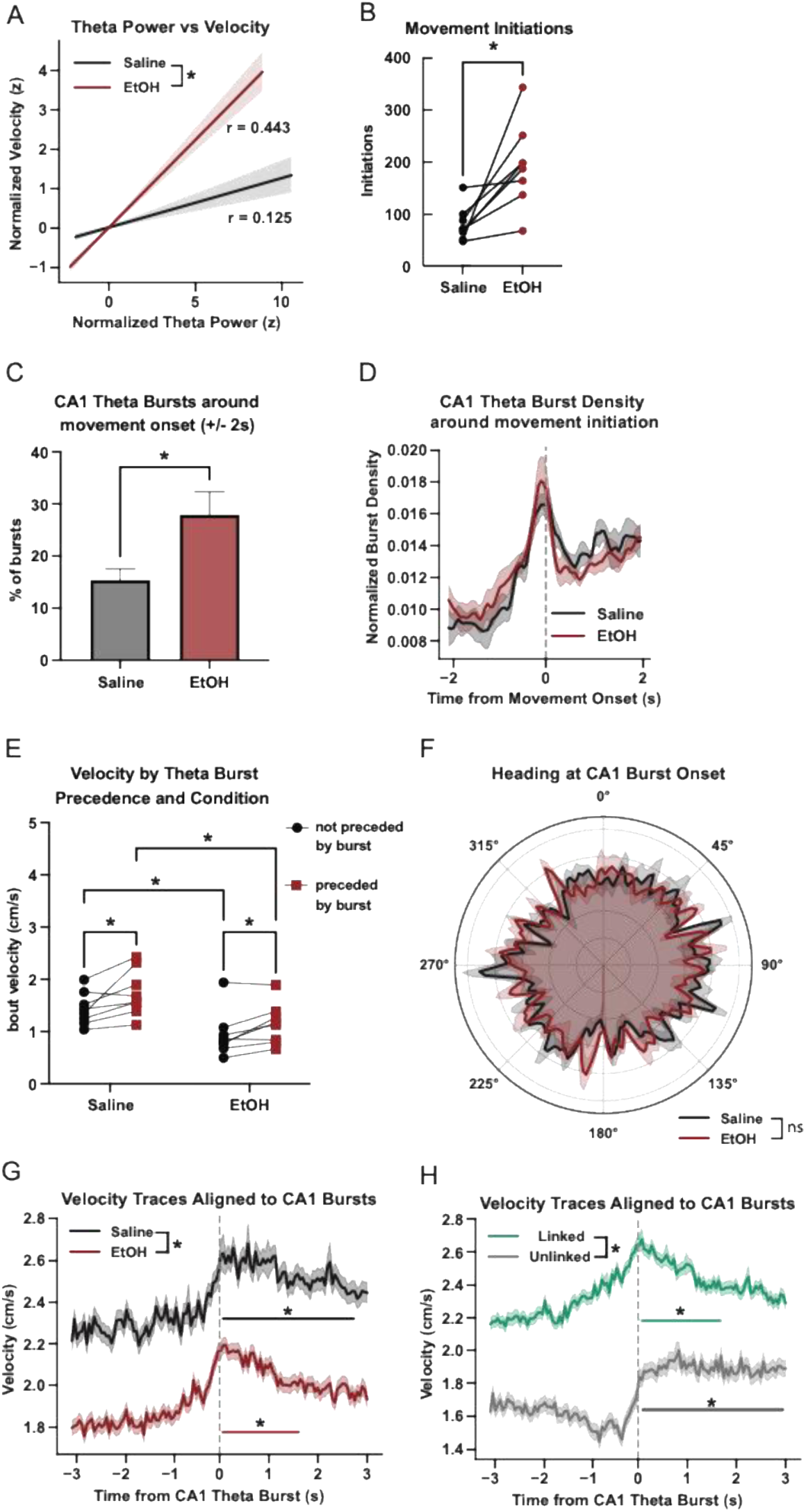
CA1 theta bursts are temporally linked to locomotor dynamics during foraging. (A) Correlation between normalized CA1 theta power and normalized velocity: saline (r = 0.125) and EtOH (r = 0.443); linear model showed a significant θ × condition interaction (p < 0.0001). (B) Movement bout initiations by treatment type (paired *t*-test, *p* = 0.0093). (C) Percentage of CA1 theta bursts within ±2 s of movement onset: Saline vs. EtOH (paired t-test, p=0.0256) (D) Group-averaged histograms show normalized burst density aligned to movement initiation (t = 0) by treatment type (EtOH: red, Saline: black). Curves represent per-mouse smoothed histograms normalized within animal and averaged by condition. (E) Comparisons of movement bout average velocity with regard to preceding burst presence (paired t-tests | Saline (preceded vs not preceded), p = 0.0006; EtOH (preceded vs not preceded), p = 0.0065, Not preceded (Saline vs EtOH), p < 0.0001; Preceded (Saline vs EtOH), p < 0.0001). (F) Polar histograms showing heading distributions at CA1 theta burst onset (Rayleigh test: Saline, p = 0.56; EtOH, p = 0.23 | MWW test: p = 0.571). (G) Mean velocity (± SEM) aligned to CA1 theta burst onset (–3 to +3 s) for Saline (black) and EtOH (red). LME with fixed effects of condition, time, and their interaction yielded: Treatment: β = 0.409, p < 0.0001; Time: β = 0.042, p < 0.0001; Interaction: β = 0.012, p < 0.0001. FDR-corrected paired t-tests against pre-burst baseline identified 45 significant post-burst bins in saline (0–2.5 s) and 23 in EtOH. Significant bins are indicated on the traces. (H) Mean velocity (± SEM) aligned to CA1 theta burst onset (–3 to +3 s) for coupled bursts (green) and uncoupled bursts (gray). LME with fixed effects of burst type, time, and their interaction yielded: Burst Type: β = –0.547, p < 0.0001; Time: β = 0.039, p < 0.0001; Interaction: β = 0.024, p < 0.0001. FDR-corrected paired t-tests against pre-burst baseline identified significant post-burst bins from 0–1.5 s for coupled bursts and from 0–3 s for uncoupled bursts. Significant bins are marked on the traces.

To assess whether theta bursts were in fact related to subsequent motor output, we quantified movement bout velocity and duration as a function of burst precedence. For each movement bout >1 second, we computed whether it was preceded by a CA1 theta burst within 2 seconds and observed that bouts preceded by a burst were consistently longer and faster than unaccompanied bouts (Fig. 5E). Post hoc comparisons confirmed that burst-preceded bouts were faster in both saline and EtOH and that EtOH increased velocity across burst and no-burst trials.

Lastly, we asked whether CA1 theta bursts exhibited directional tuning for heading. For each burst, the animal’s instantaneous heading angle was estimated as the arctangent of x and y velocity components, computed from video tracking. Circular histograms were constructed for each animal, smoothed, and averaged across mice to yield group-level polar plots. While theta bursts exhibited non-uniform heading direction in both groups, Rayleigh tests showed only weak or nonsignificant directionality and no difference in angular distributions across conditions (Fig. 5F).

To assess how CA1 theta bursts relate to movement vigor, we aligned velocity traces to burst onset (≥200 ms duration) and analyzed traces from −3 to +3 seconds around each burst. We extracted and averaged velocity across all bursts per session and condition.

A linear mixed-effects model was used to compare group-level traces, with fixed effects for condition, time, and their interaction, and a random intercept per mouse. The model revealed a significant main effect of condition, indicating overall higher velocity in saline sessions, as well as a significant time × condition interaction, reflecting a steeper increase in post-burst velocity in saline compared to EtOH (Fig. 5G). To identify specific periods of significant post-burst movement, we performed paired t-tests at each post-burst time bin (0–3 s) against the pre-burst baseline (−3 to 0 s), applying FDR correction. In saline sessions, 45 bins showed significant increases in velocity following the burst (corrected p < 0.05), spanning from 0 to ∼2.5 s. In EtOH sessions, only 23 bins were significant, indicating a shorter-lasting response. These results suggest that EtOH blunts the typical movement increase associated with hippocampal theta bursts.

We next asked whether MS activity modulated burst-related movement. Bursts were labeled as “coupled” if preceded by an MS theta burst within 1 s, and “uncoupled” otherwise. We again extracted velocity traces around each CA1 burst and grouped them by linkage. We found that uncoupled bursts were associated with lower overall velocity and a more prolonged increase in velocity over time. Post-burst binwise comparisons showed that both coupled and uncoupled bursts elicited significant increases in velocity but the duration differed: coupled bursts showed a transient increase lasting ∼1.5 s, whereas uncoupled bursts drove a more sustained increase up to 3 s post-burst (Fig 6H). We interpret these results as evidence for CA1 theta bursts relating to cognitive processes preceding movement initiation.

**Figure 6.**
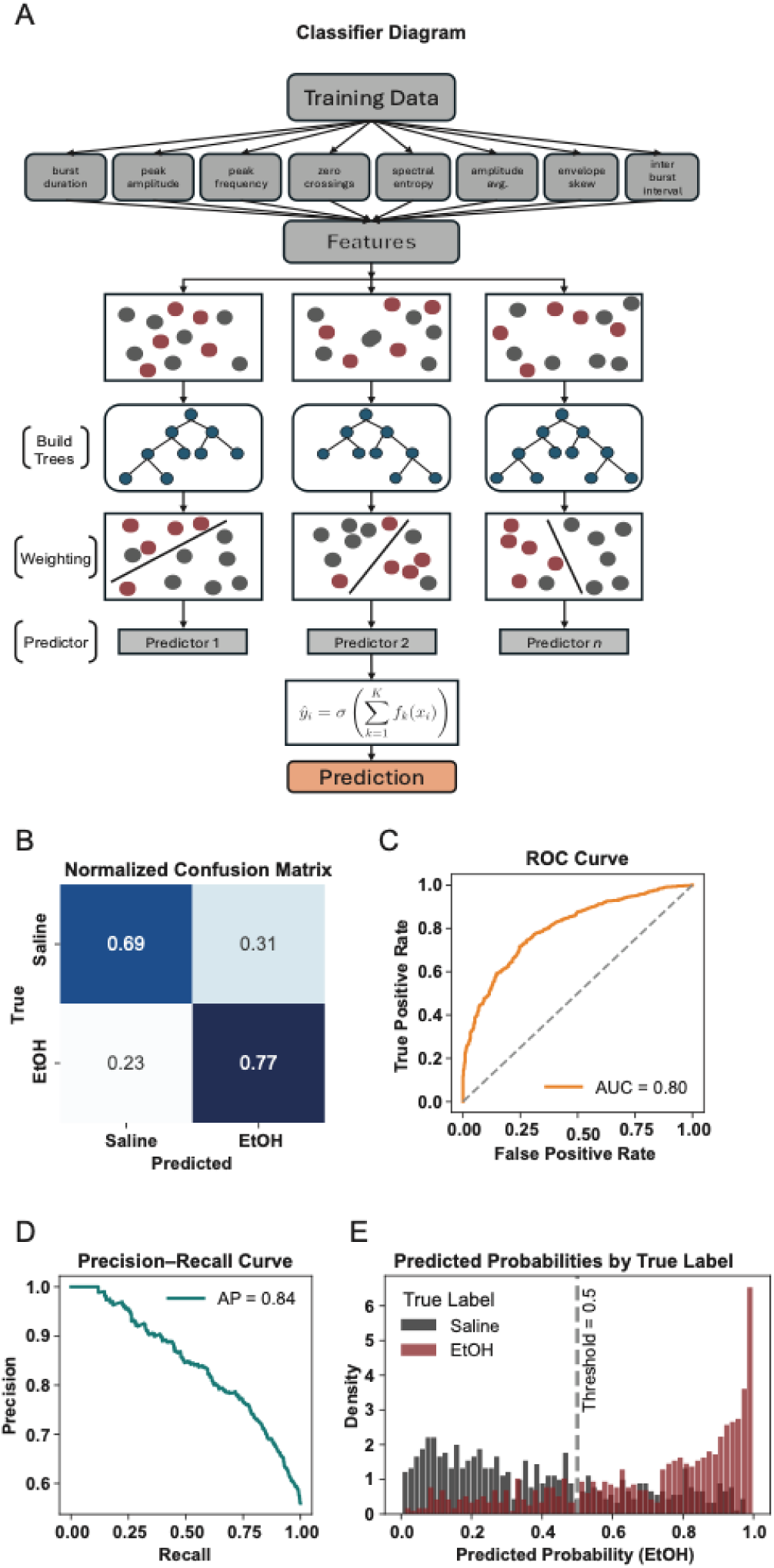
XGBoost classifier reliably distinguishes EtOH and saline conditions based on CA1 theta burst features. (A) XGBoost classification pipeline: normalized theta burst features extracted from LFP recordings are sequentially passed through an ensemble of gradient-boosted decision trees. Each tree is trained to correct the residual errors of its predecessors using a loss-minimizing split criterion. The weighted sum of tree outputs produces a logit, which is transformed via a sigmoid function into the probability of an EtOH versus saline origin. (B) Normalized confusion matrix for XGBoost burst classification (EtOH vs. saline). Overall accuracy = 74%; EtOH precision = 0.76; EtOH recall = 0.77. Misclassifications cluster near the 0.5 decision boundary. (C) ROC curve showing a high area under the curve (AUC = 0.80), indicating strong threshold-independent discriminability between conditions. (D) Precision–Recall curve for the XGBoost classifier (EtOH vs. saline), AUC = 0.82. (E) Distribution of predicted probabilities for EtOH (red) and saline (gray) bursts. Smoothed histograms show many EtOH bursts assigned probabilities > 0.8 and many saline bursts < 0.2, with the dashed vertical line marking the 0.5 decision threshold.

### Burst classification using XGBoost

Our trained XGBoost classifier (Fig. 6A) achieved strong predictive performance, with an area under the ROC curve (AUC) of 0.80 and an average precision of 0.84 (Fig. 6C– D). The normalized confusion matrix (Fig. 6B) showed an overall classification accuracy of 74%, with precision and recall values of 0.76 and 0.77, respectively, for EtOH bursts. These performance metrics confirm the model’s ability to recover EtOH-associated burst features with high reliability. Misclassifications primarily occurred near the threshold boundary, reflecting natural overlap in burst morphology between conditions. Predicted probability distributions revealed clear separation between classes, with EtOH-labeled bursts skewed toward high predicted probabilities (>0.8) and saline bursts clustering around low probabilities (<0.2) (Fig. 6E). This bimodal structure indicates the classifier was confident in a large fraction of its predictions, despite some overlap near the 0.5 decision threshold. Together, these results demonstrate that EtOH induces systematic, feature-level changes in CA1 theta bursts that can be detected by a supervised learning model. This supports the broader idea that theta bursts encode condition-specific circuit dynamics and may serve as sensitive markers of neuromodulatory state.

## Discussion

In the present manuscript we recorded LFP in three brain regions with known important roles in cognition, particularly related to theta oscillations. We report several findings related to 1) general inter-region functional connectivity and communication and 2) how a commonly used, and misused, psychoactive compound, EtOH, affects these phenomena, while mice perform a basic foraging task.

Importantly, this study provides additional evidence that theta bursts occur in the mouse hippocampus and MS [31], that these events are occur in a temporally relevant way between the two regions, and appear to correlate in time with motion initiation. Rather than broadly silencing theta activity, EtOH appears to reconfigure its temporal structure by promoting brief, high-amplitude events while weakening rhythmic continuity and degrading hippocampal stability over time. One possible mechanism by which these bursts are induced is that EtOH disrupts the intrinsic pacemaker properties of MS GABAergic neurons while simultaneously altering the excitability or entrainment of hippocampal networks. The MS contains rhythmically bursting neurons that are critical for generating hippocampal theta oscillations [12], [32] and our findings suggest EtOH may act on these pacemaker mechanisms. Increased theta burst rate but shortened burst duration in CA1 suggests that local oscillatory assemblies may be more readily triggered under EtOH, yet less able to sustain coherent activity over time. This shift may reflect changes in interneuronal recruitment or disrupted input integration from septal afferents [13], resulting in more fragmented oscillatory states consistent with known GABAergic and cholinergic modulation of theta rhythms [14].

Interestingly, EtOH preserved and in some respects enhanced short-latency MS→CA1 burst coupling. The increased probability of CA1 bursts following MS bursts, and the tighter correlation of their durations under EtOH, suggests that interregional transmission is not abolished. These findings raise the possibility that EtOH biases septohippocampal communication toward brief, tightly aligned events, potentially reflecting altered synaptic dynamics or changes in the balance between excitatory and inhibitory drive [9]. This enhanced coupling during burst events suggests a shift from continuous rhythmic coordination to episodic synchronization.

This pattern of effects—reduced stability, elevated burst rate, and sharpened MS–CA1 alignment—may represent a shift in network dynamics toward a more reactive, less integrative state. Notably, septohippocampal theta coherence was elevated during bursts and remained above pre-burst baseline after the burst concluded, indicating that these brief synchronization events have lasting effects on network coordination. Such changes may compromise the temporal binding of distributed cell assemblies that is thought to underlie memory encoding and spatial navigation [33].

The finding that low dose EtOH has detectable effects on septohippocampal theta oscillations aligns with our observations, though our focus on burst dynamics reveals more nuanced alterations than traditional spectral analyses might detect. Importantly, the current behavioral context may engage theta generators differently than classical restraint-based paradigms, offering a more translationally relevant window into EtOH’s circuit-level effects.

At a systems level, the capacity of a simple classifier to discriminate EtOH and saline sessions based on burst morphology suggests that EtOH induces consistent physiological fingerprints in hippocampal activity. Whether these signatures generalize across behavioral states or evolve with repeated exposure remains an open question, but their presence underscores the reliability of burst-level features as readouts of circuit modulation. This finding has important implications for understanding individual differences in alcohol sensitivity and developing biomarkers for alcohol-induced cognitive impairment.

Recent theoretical work has re-emphasized the role of extracellular electric fields—as captured by LFPs—not merely as epiphenomena, but as functional substrates for neural computation [34]. From this perspective, the EtOH-induced alterations we observe in theta burst morphology and coordination may reflect changes in the structure of electric fields that shape network excitability. Theta oscillations represent collective dynamics of multineuronal membrane potentials, and shorter, more frequent theta bursts could shift the spatiotemporal patterning of these fields, narrowing the window for coherent input summation and altering the firing of cell assemblies. This reframing encourages future studies to consider how pharmacological perturbations reshape field dynamics—and in turn, cognitive operations—not just through synaptic or spiking activity, but through the emergent geometry of the surrounding electrical landscape.

The preservation of burst-triggered coupling between MS and CA1 under EtOH suggests that septohippocampal communication operates through multiple temporal scales. Past experiments have shown that theta arises from interactions between medial septum and intra-hippocampal circuits, and our findings indicate that EtOH differentially affects these interaction modes. This has important implications for understanding how alcohol impairs memory formation and spatial processing.

Taken together, these findings support the idea that rapid fluctuations in theta exist, and that their dynamics are modulated by EtOH in ways that may be functionally significant. These changes may reflect neuromodulatory interference with cholinergic or GABAergic tone in MS and CA1, consistent with known pharmacological effects of EtOH on neurotransmitter systems [35]. The burst-centric nature of these alterations suggests that EtOH may particularly disrupt the temporal precision required for episodic memory formation and spatial navigation, processes that depend on sustained theta rhythmicity [36].

In sum, we report here several important features of septohippocampal communication at the level of LFP, both during normal conditions and under the influence of EtOH exposure in awake, behaving mice. Future work should test what cognitive processes these features reflect, and how changes, both natural and pharmacologically affected, translate to measurable impairments in spatial memory and behavioral flexibility, as well as how single unit activity relates to the observed LFP phenomena. Additionally, investigating dose-response relationships and the reversibility of these effects would provide important insights into the mechanisms underlying alcohol’s cognitive effects.

The development of theta burst-based biomarkers could also advance our understanding of individual vulnerability to alcohol-induced cognitive impairment.

## Author contributions

CBD and AJK designed experiments. CBD and MDM collected data. CBD and AJK performed statistical analysis of data. CBD and AJK wrote initial draft of manuscript, and after which all authors reviewed, discussed, and contributed to the final manuscript.

