## Supplemental Information for "Acute EtOH enhances septohippocampal coordination but disrupts intrinsic hippocampal theta dynamics during foraging"

### Supplemental Methods and Materials

#### Subjects

Mice were implanted at 4 months of age and were 5.5mo at the time of testing for present studies. They were group housed prior to surgery and singly housed post-operatively. Subjects were food restricted to 90% of their body weight to encourage foraging, given ad libitum access water, except during testing, and housed in an environment maintained at 22.2 °C and 50% humidity. All behavioral testing and LFP recordings took place during the first half of the light cycle. One animal's CA1 electrode placement was not correctly placed, so this animal's CA1 LFP data was not used in analysis.

##### Supplemental Methods: LFP Acquisition and Spectral Analysis

Local field potentials (LFPs) were recorded using a Tucker-Davis Technologies (TDT) PZ5 amplifier system interfaced with Synapse software, and digitized at a sampling rate of 1017 Hz. Recordings were obtained via custom-built, chronically implanted microwire electrodes and connected to the TDT system through Intan RHD headstages. Signals from three brain regions—medial septum (MS)**,** dorsal hippocampus (CA1), and medial prefrontal cortex (mPFC)**—**were continuously acquired during 20-minute behavioral sessions. A reference electrode was implanted over the cerebellar plate.

Raw LFP signals were preprocessed using custom Python scripts. To remove ambient electrical noise, signals were first notch filtered at 60 Hz using a zero-phase bandstop FIR filter implemented via the NeuroDSP library (filter_signal, pass_type='bandstop', f_range=(57–63 Hz), n_cycles=8). This provided sharp stopband attenuation while preserving phase fidelity. Following line noise removal, signals were bandpass filtered into canonical frequency ranges using zero-phase finite impulse response (FIR) filters with narrow transition bands and minimal phase distortion. The frequency bands used were: Delta: 0.5–4 Hz, Theta: 5–12 Hz, Beta: 13–20 Hz, Low Gamma: 30–55 Hz, and High Gamma: 65–100 Hz. Filters were designed per-band using the firwin method with custom settings for number of cycles and maximum duration, ensuring high stopband attenuation and temporal precision. All filtering steps were applied using forward–reverse (filtfilt) convolution to achieve zero-phase distortion.

For spectral analysis, power spectral density (PSD) was estimated using Welch's method, implemented in the compute_spectrum() function from the NeuroDSP package. LFP signals were segmented into 6-second non-overlapping windows, and the PSD of each window was computed with a Hann window and nfft = 8096, yielding high frequency resolution up to ~140 Hz. PSD estimates were converted to relative power by dividing the absolute power in each frequency band by the total power from 0–55 Hz, effectively normalizing spectral content across sessions and animals:

This approach enabled comparison of oscillatory dynamics across conditions independent of total signal energy. Relative power values were computed for each region and frequency band across all sessions and time windows and stored in structured dataframes for downstream statistical analysis and visualization.

All signal processing, including filtering, PSD estimation, and relative power computation, was performed using a modular Python pipeline incorporating NeuroDSP, SciPy, and custom scripts to ensure reproducibility and cross-session consistency. Note this filtering procedure was only used for PSD calculation.

#### Burst Detection

To isolate transient theta-band activity for burst-level analysis, we employed the following protocols.

*Preprocessing and Filtering:* Raw LFP signals were recorded at a sampling rate of 1017 Hz and notch-filtered at 60 Hz to remove electrical line noise. Signals were then bandpass filtered in the theta range using zero-phase finite impulse response (FIR) filters. Specifically, a 5–12 Hz passband was applied using the filter_signal() function from the NeuroDSP package, which preserves phase relationships and minimizes temporal distortion. Filter parameters were set using 3–4 cycles of the low cutoff frequency, with duration capped to prevent edge artifacts. Edge samples affected by filtering were removed to ensure clean analytic estimation.

*Envelope Computation and Normalization:* For each filtered signal, we computed the analytic amplitude envelope using the magnitude of the Hilbert transform. The resulting envelope was z-scored relative to the entire session to normalize across animals and sessions and to enable amplitude thresholding based on standard deviations from the mean.

*Dual-Threshold Burst Detection:* Theta bursts were identified using a dual-threshold crossing algorithm applied to the z-scored envelope: Burst onset was defined as the point where the envelope exceeded a high threshold of $z > 2.0$, the burst continued as long as the envelope remained above a lower threshold of $z > 1.0$, burst offset was defined as the point where the envelope dropped back below the low threshold, bursts were retained only if their duration exceeded a minimum threshold of 100 ms. This method allows for robust detection of high-amplitude theta epochs while avoiding fragmentation of longer bursts and suppressing spurious, short excursions (see Supplemental Figure 1).

*Duration Constraint Based on Oscillatory Cycle Count:* To ensure that retained bursts reflect true oscillatory episodes rather than high-amplitude noise or artifacts, we imposed a post-hoc duration filter based on the lower frequency bound of the theta band (5 Hz). Specifically, we required that each burst exceed 1.5 cycles of the lowest frequency, corresponding to a minimum duration of ≥ 300 ms (i.e., 1.5 × (1/5) seconds). This constraint was applied after initial detection to exclude events inconsistent with known theta periodicity.

#### MS–CA1 Phase Offset and PLV Analyses

To assess alcohol's impact on septohippocampal timing, we analyzed instantaneous phase relationships between medial septum (MS) and dorsal hippocampus (CA1) local field potentials (LFPs) within the theta frequency band (5–12 Hz). This analysis was performed on a per-sample basis using Hilbert transform-derived phase angles from bandpass-filtered LFP signals.

Signal Preparation and Phase Extraction
LFPs were bandpass filtered in the theta range using zero-phase FIR filters designed with 4–8 cycles of the lower cutoff frequency. Filter coefficients were computed using firwin and applied using forward convolution (lfilter). The analytic signal for each filtered trace was then computed using the Hilbert transform, and the instantaneous phase was extracted as the angle of the complex-valued result.

For each session, the phase offset between MS and CA1 was computed as:

$$\Delta\phi(t) = \mathrm{angle}\left(e^{i\left(\varphi_{\mathrm{MS}}(t) - \varphi_{\mathrm{CA1}}(t)\right)}\right)$$

These offsets were wrapped to the range [−π, π], producing a per-sample distribution of phase differences per mouse, session, and condition.

*Phase Offset Distribution and KDE Analysis:* To compare conditions, circular histograms of MS–CA1 phase offsets were computed per mouse for each treatment group (SALINE, ALC) and smoothed using a Gaussian kernel (σ = 2). Resultant kernel density estimates (KDEs) were visualized using polar plots and aggregated across mice. Differences in KDEs between treatment groups were assessed using a permutation-based L2 norm comparison and the Mardia-Watson-Wheeler circular ANOVA, testing for both shape and centrality differences.

*Burst-Aligned Phase Analysis:* To probe how MS–CA1 phase relationships varied around CA1 theta bursts, we compared bursts that were preceded by an MS theta burst within 200 ms ("coupled") against those without recent MS activity ("uncoupled"). Bursts were identified using a dual-threshold algorithm (see Burst Detection Methods), and burst timing metadata were used to segment windows around each CA1 burst (1250 ms post-burst window). MS and CA1 signals from these windows were filtered and phase-offsets computed as above. The circular mean of phase differences per burst was used for analysis.

KDEs were generated per mouse per condition and grouped by coupling status. Group-level comparisons of preferred phase angles and mean resultant vector lengths (MRVLs) were visualized via polar arrows and evaluated statistically using the Mardia-

Watson-Wheeler test. Additional comparisons of coupled vs. uncoupled distributions were performed using permutation tests across angle bins.

*Theta PLV Between CA1 and MS During Coupled Events:* To assess cross-regional coordination beyond MS–CA1, we examined theta-band phase-locking values (PLV) between CA1 and MS around coupled theta bursts. Coupled bursts were defined as above. For each event, 2-second windows before and after the CA1 burst were extracted from bandpass-filtered signals, and PLV was computed as:

$$\mathrm{PLV} = \left| \frac{1}{N} \sum_{t=1}^{N} e^{i\left(\varphi_{\mathrm{CA1}}(t) - \varphi_{\mathrm{MS}}(t)\right)} \right|$$

Pre- and post-burst PLVs were computed per event, and the PLV ratio (post/pre) was averaged per mouse to assess event-related changes in CA1–MS coordination under saline and EtOH conditions.

#### Lag Estimation Between MS and CA1 Theta

To quantify the temporal delay between MS and CA1 theta oscillations, we computed cross-correlation lags between their respective amplitude envelopes. LFPs were bandpass filtered in the theta range (5–12 Hz) using zero-phase FIR filters. The analytic amplitude (envelope) was obtained by applying the Hilbert transform to each filtered signal. Envelopes were z-scored to normalize across sessions.

Each session was segmented into overlapping 8-second windows with a 2-second step size**.** Within each window, the cross-correlation was calculated between the MS and CA1 z-scored amplitude envelopes. We restricted the lag search to a window of ±150 milliseconds (±153 samples at 1017 Hz). The lag corresponding to the peak cross-correlation within this range was recorded as the estimated MS→CA1 delay for that window.

For each session, we computed the mean lag as well as three measures of lag variability: standard deviation (SD)**,** interquartile range (IQR)**,** and median absolute deviation (MAD). To assess directional differences across treatment conditions, lag metrics were averaged per mouse and compared between saline and ethanol sessions using paired t-tests**.**

To visualize within-mouse changes in lag distribution, we subtracted the session median from each lag (centering), and generated normalized density histograms across all mice. This approach emphasizes relative changes in timing precision independent of absolute delays.

#### XGBoost Classification of CA1 Theta Bursts

To assess whether CA1 theta burst characteristics reliably distinguished ethanol (EtOH) exposure from saline, we implemented a supervised machine learning pipeline using XGBoost. Theta bursts were detected using a dual-threshold algorithm and retained if they lasted ≥200 milliseconds, ensuring physiological plausibility for theta-band activity.

Feature Extraction

For each burst, we extracted nine numerical features describing its shape and spectral content:

1. Duration (ms)
2. Peak amplitude
3. Mean amplitude
4. Zero-crossing count
5. Peak frequency (from multitaper power spectral density within the 5–12 Hz range)
6. Envelope skewness (computed from the analytic signal)
7. Normalized spectral entropy (estimated from the theta-band PSD)
8. Inter-burst interval (IBI, ms) — time since the previous CA1 theta burst
9. MS–CA1 latency (ms) — time since the last MS theta burst ended before the CA1 burst

The MS–CA1 latency feature was set to NaN for bursts not preceded by an MS burst. All features were scaled using a MinMaxScaler, and missing values were imputed using column medians.

To account for between-subject variability, mouse identity was one-hot encoded and concatenated to the feature matrix. This enabled the model to consider individual baseline differences without leaking session condition information.

Model Training and Evaluation

The dataset was split into stratified training (80 percent) and test (20 percent) sets, maintaining class balance. We trained an XGBoost classifier with the following parameters: n_estimators = 300, max_depth = 10, and learning_rate = 0.05.

The model's performance was assessed using the confusion matrix, receiver operating characteristic (ROC) curve and area under the curve (AUC), and precision–recall (PR) curve with average precision (AP).

All analyses were conducted in Python (v3.11) using the XGBoost, scikit-learn, NumPy, NeuroDSP, and Pandas libraries.

### Supplemental Figures

#### Supplemental Figure 1

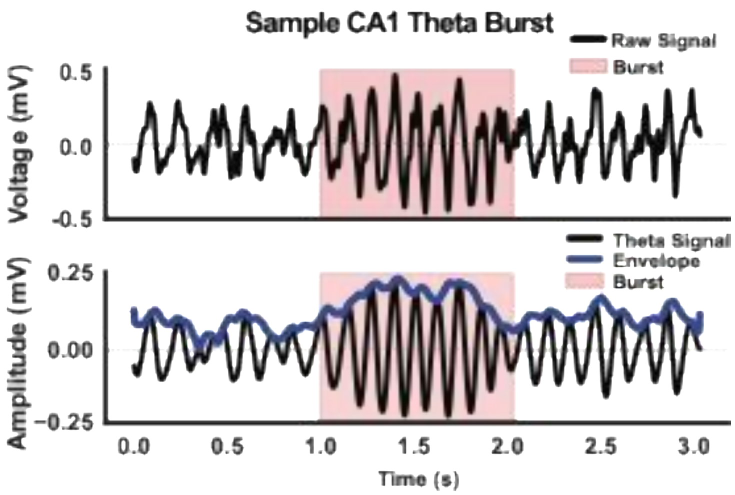


**Sample CA1 theta burst.**

(Top) The raw LFP trace (1–100 Hz) from CA1 is shown with the detected burst region highlighted in red. (Bottom) The theta-filtered signal (5–12 Hz, black) and its Hilbert envelope (blue) illustrate amplitude dynamics used for burst detection.

#### Supplemental Figure 2

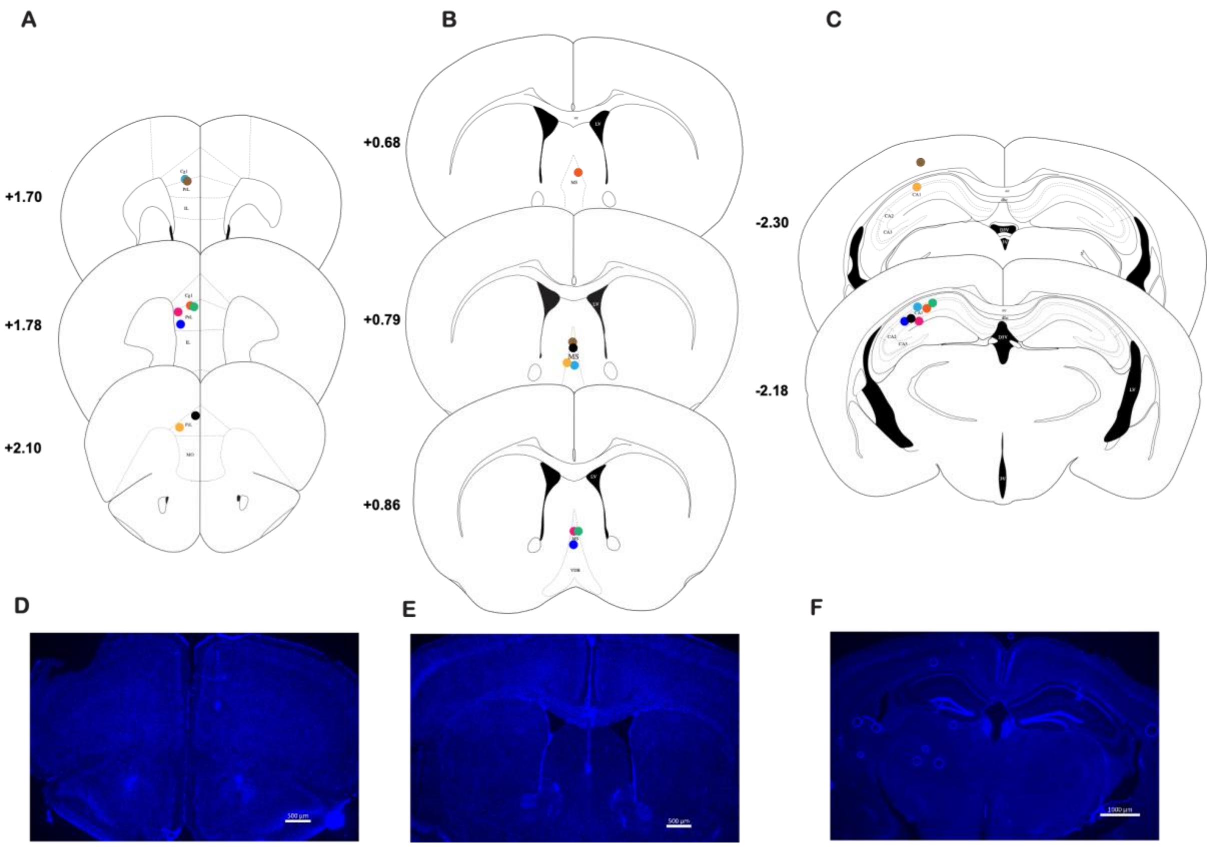


##### Electrode placements.

(A–C) Schematic coronal sections illustrating electrode tip locations for all mice (n = 8). Each color represents a different animal. Electrodes targeted the medial prefrontal cortex (mPFC; A), medial septum (MS; B), and dorsal CA1 of the hippocampus (C). Numbers indicate anterior-posterior distance from bregma.

(D–F) Representative post-mortem histological sections from one animal showing final electrode placement sites. Electrolytic lesions were made at the end of the experiment to mark electrode tips. Blue indicates DAPI nuclear stain. Regions shown: mPFC (D), MS (E), and CA1 (F). D,E scale bar = 500uM, F scale bar = 1000uM.
